# Next Generation-Targeted Amplicon Sequencing (NG-TAS): An optimised protocol and computational pipeline for cost-effective profiling of circulating tumour DNA

**DOI:** 10.1101/366534

**Authors:** Meiling Gao, Maurizio Callari, Emma Beddowes, Stephen-John Sammut, Marta Grzelak, Heather Biggs, Linda Jones, Abdelhamid Boumertit, Sabine C Linn, Javier Cortes, Mafalda Oliveira, Richard Baird, Suet-Feung Chin, Carlos Caldas

## Abstract

Circulating tumour DNA (ctDNA) detection and monitoring has enormous potential clinical utility in oncology. We describe here a fast, flexible and cost-effective method to profile multiple genes simultaneously in low input cell-free DNA (cfDNA): Next Generation-Targeted Amplicon Sequencing (NG-TAS). We designed a panel of 377 amplicons spanning 20 cancer genes and tested the NG-TAS pipeline using cell-free DNA from two hapmap lymphoblastoid cell lines. NG-TAS consistently detected mutations in cfDNA when mutation allele fraction was >1%. We applied NG-TAS to a clinical cohort of metastatic breast cancer patients, demonstrating its potential in monitoring the disease. The computational pipeline is available at: https://github.com/cclab-brca/NGTAS_pipeline.

## BACKGROUND

Cell free DNA (cfDNA) in plasma was first analysed in cancer patients nearly 50 years ago [1]. A fraction of cfDNA was shown to carry mutations found in the matched tumour, and designated circulating tumour DNA (ctDNA) [1–3]. The utility of ctDNA as a non-invasive diagnostic, prognostic or predictive biomarker in human cancer is now well documented [4–8].

The amount of cfDNA in plasma is usually low and the ctDNA fraction is typically only 1-30%, hence low mutant allele frequencies have to be detected. Human cancers are genetically heterogeneous and mutations occur infrequently at recurrent hotspots. Therefore, in most clinical scenarios (e.g. early diagnosis or monitoring of tumour evolution), high sensitivity and the simultaneous investigation of multiple gene targets are desirable features of any ctDNA detection and quantitation method.

There are a range of methods for detecting mutations in ctDNA, with the target varying from a single nucleotide variant (SNV) to the whole genome. A widely used method to detect mutations in ctDNA is digital polymerase chain reaction (dPCR) performed in microfluidic devices or water-in-oil droplet emulsions [9,10]. Whilst dPCR is able to detect rare mutations with extremely high sensitivity, it is restricted by the number of targets that can be examined in a single reaction [11].

Several sequencing based approaches have been developed to incorporate multiple genomic loci, enabling *de novo* mutation identification in ctDNA. Previously, we described Targeted Amplicon Sequencing (TAm-Seq), which utilised 48 primer pairs to identify mutations in hotspots or selected regions of 6 key driver genes [12]. While TAm-Seq is useful, it is limited to a small number of targets. Capture based sequencing methods can cover a larger number of genes (or the whole exome), but are costly at the sequencing coverage (>300) required to detect allele frequencies ~1%.

There several ready to use commercial kits for ctDNA sequencing, which can cover up to hundreds of mutation hotspots and many genes. These include Invision™ (Inivata), Oncomine™ cfDNA assay (Thermo Fisher Scientific), Guardant360^TM^ (Guardant Health), and PlasmaSELECT™ (Personal Genome Diagnostics). These products are expensive and test custom gene panels. Disturbingly, a recent study comparing the performance of two of these commercial products (Guardant360^TM^ and PlasmaSELECT™) in a cohort of plasma samples from prostate cancer patients, revealed poor agreement [13].

Recently unique molecular barcodes have been developed to tag each cfDNA template molecule before PCR amplification in order to reduce the error rate and allow robust detection of rare mutant alleles in ctDNA [14].

In summary, using current ctDNA profiling methodology, the detection of mutations in a good number of cancer genes with sufficient sensitivity and in a cost-effective way poses significant challenges. Here we describe a new method for the profiling of ctDNA, designated Next Generation-Targeted Amplicon Sequencing (NG-TAS), with several unique features: i) optimised for low input ctDNA; ii) high level of multiplexing, enabling the analyses of multiple gene targets; iii) a bespoke computational pipeline for data analysis; and iv) very competitive costing. NG-TAS is designed to be flexible in terms of choice of gene targets and regions of interest; thus, it can be tailored to various cancer types and clinical contexts.

## METHODS

### Patient samples and blood processing

Patients were recruited from 3 different centres including Cambridge University Hospital, Netherland Cancer Institute (NKI) and Vall d’Hebron Institute of Oncology (VHIO). Metastatic breast cancer patients with hormone receptor positive tumours were recruited as a part of a clinical trial (patient number = 30, plasma samples number = 366). Blood samples were collected in EDTA tubes and processed within one hour to prevent lymphocyte lysis and fragmentation. Samples were centrifuged at 820g for 10min at room temperature to separate the plasma from the peripheral blood cells. The plasma was further centrifuged at 1400g for 10min to remove remaining cells and cell debris. The plasma was stored at −80°C until DNA extraction. This study was approved by the regulatory and ethic committees at each site and the reference number is NCT02285179 (https://clinicaltrials.gov/ct2/show/NCT02285179). All human samples used were collected after informed consent, and the study was fully compliant with the Helsinki Declaration

### DNA extraction from plasma and buffy coat

Plasma DNA was extracted from between 2-4 ml of plasma with the QiaSymphony according to the manufacturer’s instruction using Qiagen circulating DNA extraction kit. DNA was isolated from the buffy coat samples using DNeasy Blood & Tissue Kits (Qiagen).

### Generation of cfDNA from NA12878 and NA11840

As previously reported [15] two lymphoblastoid cell lines, NA12878 and NA11840 from Human Genome Diversity Project (HGDP)-CEPH collection were obtained from the Coriell Cell Repository. A catalogue of highly accurate whole genome variant calls and homozygous reference calls has been derived for sample NA12878 by integrating independent sequencing data and the results of multiple pipelines (http://www.illumina.com/platinumgenomes). NA11840 cell line was chosen from a set of 17 available CEPH cell lines in our laboratory as it shared the least number of SNPs with NA12878, to generate the maximum number of virtual somatic SNVs.

The cell lines were grown as suspension in RPMI 1640-Glutamax (Invitrogen) supplemented with 10% foetal calf serum, 5% penicillin and streptomycin at 37°C and 5% CO_2_. The media that the cell lines were grown in were collected when cells were passaged. The media were centrifuged at 1500rpm for 10 minutes at 4°C to remove cells and cellular debris. The clarified media were stored at −20°C until required. Cell-free DNA was extracted from the thawed media using the Qiagen circulating DNA extraction kit (Qiagen) according to the manufacturer’s instructions and quantified using Qubit High Sensitivity DNA quantification kit (Life Technologies). DNA from both cell lines was diluted to obtain 50 ng/μl stock concentrations. To generate the serial dilutions of one cell line with the other, we mixed by volume to obtain the percentage (volume/volume) as presented in Additional file 1: Table S1 (n=12).

Platinum variant calls for sample NA12878 (the virtual ‘tumour’) and confident regions (high confidence homozygous reference regions plus platinum calls) [16] were downloaded from http://www.illumina.com/platinumgenomes. Genotype data for sample NA11840 (the virtual ‘normal’) was obtained from the 1000 Genomes website. Platinum calls were intersected with our NG-TAS panel target regions and variants shared with the NA11840 sample were excluded. Five platinum calls were covered theoretically by our NG-TAS panel; however, one was targeted by one of the amplicons showing no coverage, therefore 4 SNVs were considered as identifiable ‘somatic variants’.

### NGS library construction

NGS libraries were prepared from 3-5 ng of cfDNA using the ThruPLEX® Plasma-seq kit (Rubicon Genomics, USA) as described in the manufacturer’s instructions. NGS library was quantified using qPCR KAPA Library Quantification kit (KAPA Biosystem), while the fragment size and the NGS library yield were measured with 2200 TapeStation instrument (Agilent).

### Digital PCR

BioMark system from Fluidigm has been used for dPCR, and the analyses have been performed as previously described [17]. As described in manufacturer’s instructions, DNA samples were mixed with 2X TaqMan® Gene Expression Master Mix (Life Technology, 4369016), 20X GE Sample Loading Reagent (Fluidigm, 85000746) and 20X gene-specific assays. The reaction mix were loaded on the qdPCR 37K^TM^ IFC (Fluidigm, 100-6152). For *KRAS* (*G13D*) and *AKT1* (*E17K*) mutant and wild type PrimePCR™ ddPCR™ Mutation Assays were obtained from Bio-Rad (dHsaCP2000013 and dHsaCP2000014, dHsaCP2000032 and dHsaCP2000031 respectively). The *PIK3CA* and *ESR1* probes and primers were previously described [7,18], and the primer and probes used are listed in Additional file 1: Table S2.

### NG-TAS protocol

#### Primer design for NG-TAS

Primers were designed with NCBI Primer-BLAST tool with Tm range of 59-61°C. The universal primer sequences (CS1 and CS2) were added at the 5’ end of the designed primers. All primer pairs were tested alone and in multiplexed PCR reactions using 10 ng of TaqMan® Control Human Genomic DNA (Thermo Fisher Scientific) in 10 μl reaction volumes. The coverage and performance of primers were analysed using 2200 TapeStation instrument (Agilent) and Hi-seq 4000. The primers were grouped together as 7-8plex, and primers in each group were chosen to target different genes in order to minimise non-specific amplification and cross-reactivity.

#### Access Array™ microfluidic system

The 377 pairs of optimised primers were divided into 48 wells, with each well containing 7-8 pairs of primers for multiplexed PCR. Primers were diluted to the final concentration of 1 μM to make 20X primer solution. 4μl of the 20X primer solution from the 48 wells were added to the primer inlets of the Access Array™ IFC (Fluidigm). For the sample inlets, pre-sample master mix consisted of 2X Master Mix (Qiagen, 206143), 5X Q-solution, 20X Access Array™ Loading Reagent (Fluidigm), and DNA sample were added. The loaded IFC then moved to FC1™ Cycler (Fluidigm) for thermal cycles: 95°C for 15min, 30 cycles of 94°C for 30sec, 59°C for 90sec, 72°C for 90sec, and final extension step 60°C for 30min. The reaction products were harvested using Post-PCR IFC controller as described in manufacturer’s instructions.

The harvested product was diluted (1:10) with water for further barcoding PCR. Barcoding PCR reaction master mix contains 2X Master Mix (Qiagen), diluted harvested product from Access Array™, and Access Array™ Barcode Library for Illumina® Sequencers single direction for barcoding primers (Fluidigm, 100-4876). The thermal cycle for barcoding is: 95°C for 10min, 15 cycles of 95°C for 15sec, 60°C for 30sec, 72°C for 1min, and final extension step of 72°C for 3min. The PCR reaction was performed using T100™ Thermal Cycler (Bio-Rad).

#### Quantification and clean-up of barcode Access Array™ harvest

After barcoding PCR, all samples were analysed using 2200 TapeStation (Agilent) to measure the concentration and size of the products (average 260bp). The PCR products were pooled and cleaned with AMPure XP beads (Beckman Coulter, A63880) following the manufacturer’s instruction. Briefly, the samples were mixed with the magnetic beads to the ratio of 180:100 in volume. The beads were washed twice with 80% ethanol and dried by incubating at 30°C for 10min. Then the beads were eluted with water and the cleaned PCR product was run on the E-Gel® 2% agarose gel (Thermo Fisher Scientific, G501802) for further size selection and extraction. The band between 200-300bp was cut out and DNA was isolated from the gel using the QIAquick Gel Extraction kit (Qiagen, 28704), and 10-20nM of the eluents was submitted for paired-end Hi-seq 4000 for sequencing.

### Analysis of NG-TAS data

#### Quality control, alignment and bam files annotation

For each sequencing lane, quality control of raw data was performed using FastQC (http://www.bioinformatics.babraham.ac.uk/projects/fastqc/). Up to 384 samples were multiplexed in a single sequencing lane and demultiplexing was performed using inhouse software.

Alignment, read trimming (at 80bp) and base quality recalibration was performed in a single step using Novoalign (v 3.08). However, to facilitate a broad use of the pipeline, a version using BWA-MEM is also available. Alignment and bam metrics were computed using Picard Tools (v 2.17). To remove potential off-target PCR products, only reads mapped in proper pair and with insert size > 60bp were retained. After this filtering, bam files were locally realigned using the Genome Analysis Toolkit (GATK, v 3.6). Reads were then assigned to the amplicon they belonged to using a custom java script, in order to enable a per amplicon coverage and mutation calling analysis. Coverage was computed for each amplicon in each sample using a custom java/R script. One amplicon (SF3B1_D0069_001) showed an extremely high rate of mismatches and indels in all the analysed samples, therefore we excluded it from downstream analyses.

#### Mutation calling

Mutation calling was run separately for each amplicon in the panel. The core mutation calling was performed for each pair of plasma and normal samples (or NA12878 an NA11849 from the dilution series) using Mutect2 (included in GATK 3.6). The *minPruning* parameter was set at 5 to reduce computational time with no significant impact on the results. Besides the set of mutations passing all internal filters, we included those failing the following internal filters or a combination of them: “alt_allelejn_normal”, “clustered_events”, “homologous_mapping_event”, “multi_event_alt_allelejn_normal”. On this set of candidate mutations, we applied the following filtering criteria: coverage in normal and plasma >100x, alternative allele in normal <1% and plasma/normal VAF ratio >5. The core mutation calling was repeated for the three replicates generated for each pair and only mutations called in at least two replicates were retained. For this set of mutations, we run HaplotypeCaller (included in GATK 3.6) to compute the average VAF across the three replicates and filter out mutations with an average VAF<1% and an average plasma/normal ratio <5 (Figure 4A). An extra filter was introduced for FFPE samples, where C>T and G>A transitions with VAF<15% were filtered out because likely to be consequence of cytosine deamination caused by fixation.

In calling somatic mutations from set of longitudinal samples from the same patient, we first repeated the above procedure for all samples. Then, HaplotypeCaller was run again to estimate in all samples the coverage and VAF of each mutation called in at least one of them. This was followed by a variant annotation step using Annovar. Finally, results obtained for all amplicons were merged to generate a single vcf file. A final filter was applied at group level, that is, keeping only mutations that at least in one sample had VAF higher than 5% (Additional file 1: Fig. S1).

## RESULTS

### Optimising targeted deep sequencing in cfDNA by NG-TAS

We designed 377 pairs of primers covering all exons or hotspots of 20 genes commonly mutated in breast cancer (Table 1). To identify the genes or hotspots of interest, we primarily looked at the comprehensive study carried out in our lab (Pereira et al. Nat Comm 2016). Other genes genes (e.g. *ESR1*) were included because reported as frequently mutated in metastasis [19]. Since the average cfDNA fragment size is 160-170bp, NG-TAS primers were designed to generate amplicons of 69-157bp (Additional file 2).

**Table 1.** List of genes and regions covered in the panel.

| Gene | Target region | Hotspot position | No of ampicons |
| --- | --- | --- | --- |
| *AKT1* | Hotspot | E17<br>AA23 – 59<br>AA65 - 94 | 4 |
| *BRAF* | Hotspot | V600 | 1 |
| *Her2* | Hotspot | S310<br>AA428 - 438<br>AA746 – 797<br>AA832 – 986 | 14 |
| *HRAS* | Hotspot | AA3 – 35 (G12 and<br>G13)<br>AA49 – 77 (Q61 and<br>A66) | 3 |
| *IDH2* | Hotspot | AA132 - 162 | 1 |
| *KRAS* | Hotspot | G12 | 1 |
| *SF3B1* | Hotspot | K700 | 1 |
| *ESR1* | Part of exons | Exon 8 - 10 (LBD) | 10 |
| *SMAD4* | Part of exons | Exon 8 – 12 | 10 |
| *CDH1* | All exons |  | 46 |
| *CDKN1B* | All exons |  | 9 |
| *FOXA1* | All exons |  | 18 |
| *GATA3* | All exons |  | 23 |
| *MAP2K4* | All exons |  | 22 |
| *MAP3K1* | All exons |  | 75 |
| *PIK3CA* | All exons |  | 59 |
| *PIK3R1* | All exons |  | 11 |
| *PTEN* | All exons |  | 24 |
| *RUNX1* | All exons |  | 24 |
| *TP53* | All exons |  | 21 |

In a preliminary optimization step, individual primer pairs were tested in conventional single and multiplexed (7-8plex) PCR reactions. The NG-TAS experimental workflow (Figure 1A), starts with a multiplexed PCR step (7-8 primer pairs) performed using Access Array™, a microfluidic system from Fluidigm. Each multiplexed reaction contained primers targeting different genes to minimise the generation of unwanted PCR products. The multiplexed PCR products were assessed using the Bioanalyser and 2200 TapeStation instrument (Agilent Genomics; Additional file 1: Fig. S2). Multiplexed PCR products were then pooled and barcoded with 384 unique barcodes in a second PCR reaction. Barcoded products were pooled, and size selected to remove primer dimers before submission for NGS paired-end 150bp sequencing.

**Figure 1.**
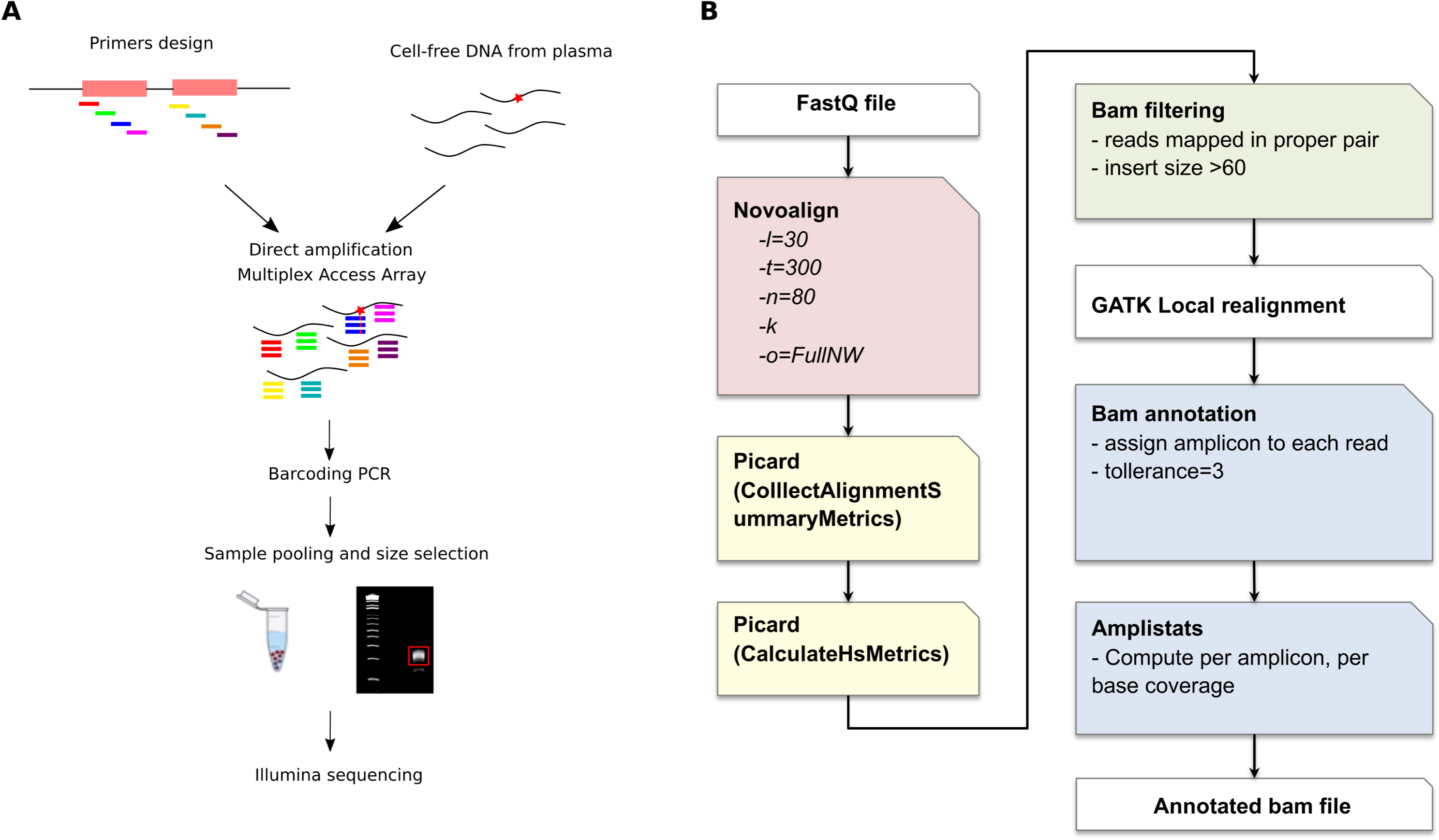
NG-TAS workflow and alignment pipeline. (A) NG-TAS workflow. Primers were designed and multiplexed for direct amplification in cfDNA obtained from plasma using Fluidigm Access Array™. The PCR products were harvested and barcoded in a subsequent PCR reaction, the samples were pooled and size selected for sequencing on an Illumina Hi-Seq 4000. (B) Schematic representation of the computational pipeline for reads alignment, filtering and annotation.

Raw sequencing data were aligned and processed as described in Figure 1B and Methods. Specific filters were applied to exclude reads from primer dimers or other PCR artefacts. Since the amplicons are partially overlapping, each read was assigned to its respective amplicon, to enable a per-amplicon analysis for coverage estimation and mutation calling.

To optimise NG-TAS we used cfDNA isolated from the culture media of the Platinum Genome Hapmap NA12878 cell line. The size profile of cfDNA isolated from the tissue culture media was similar to that of plasma cfDNA (Additional file 1: Fig. S3). We tested a range of input cfDNA amounts with NG-TAS (0.016 ng to 50 ng) in 4 replicates for each input. For each cfDNA input we tested: i) a pre-amplification step and ii) the use of the Qiagen Q solution. To assess the data generated the percentage of aligned sequencing reads was computed (Figure 2A). In the TAM-Seq protocol addition of a pre-amplification step reduced the probability of nonspecific amplification and biased coverage [12]. However, using NG-TAS the pre-amplification step reduced the percentage of aligned reads in all cfDNA input samples tested. Hence, we eliminated pre-amplification from the NG-TAS protocol. Adding Q solution systematically increased the percentage of aligned reads, with the largest improvement observed with 0.4 and 2 ng input samples (Figure 2A). Thus, we incorporated the Q solution in all subsequent NG-TAS experiments.

**Figure 2.**
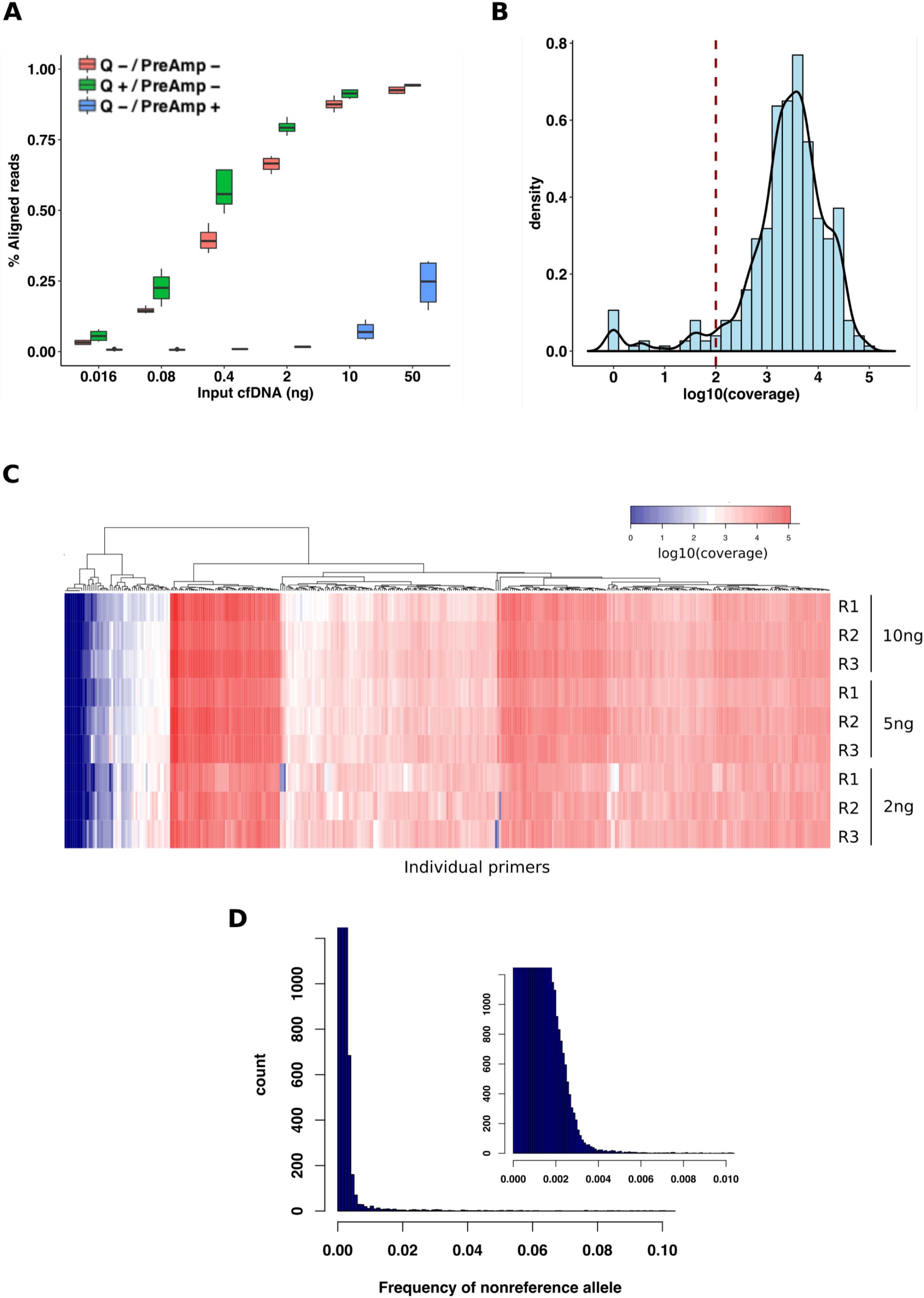
Optimising targeted deep sequencing by NG-TAS. (A) Percentage of aligned reads were compared in different samples where a variable amount of input control genomic DNA was used (range 50 to 0.016ng). The effect of pre-amplification and Q solutions are shown, red = No Q solution and no preamplification step, green = With Q solution and no pre-amplification, blue = No Q solution and with pre-amplification. (B) Density plot showing the log10 coverage values for all primers in the 10 ng NA12878 cfDNA sample. The dotted line indicates 100x coverage; median value for the distribution is 3064x. (C) Coverage heatmap of individual primers for different amount of input NA12878 cfDNA. For each amount of input DNA the analysis was performed in triplicate. (D) Distribution of all non-reference base frequencies across all target regions in the NA12878 dilution series in (C); the smaller plot on the right is a magnification of the main plot between 0 and 0.01.

We then used the optimised NG-TAS protocol in triplicate experiments for each input NA12878 cfDNA (2 ng, 5 ng, and 10 ng). With 10ng of input cfDNA NG-TAS generated a median read depth of 3064x, and only 22/377 amplicons (5.8%) had coverage less than 100x (Figure 2B). In fact, high amplicon coverage was observed irrespective of amount of input cfDNA (Additional file 1: Fig. S4A and S4B). The coverage heatmap of individual amplicons showed similar patterns with 10 ng and 5 ng cfDNA input. Strong consistency was observed within each triplicate (Figure 2C). However, with 2ng cfDNA input we observed stochastic reduction in coverage for some of the amplicons. This is probably due to a reduction in template availability, with the number of amplifiable copies approaching zero for some of the amplicons.

Using these data, the background noise was estimated by computing the average frequency for non-reference bases in each position, and for 99% of the targeted genomic positions background noise was ≤0.3% (Figure 2D).

### Sensitivity and specificity of mutation detection in control cfDNA

To establish an analysis pipeline and assess the performance of NG-TAS, we generated a benchmark dilution series, similar to what we have previously described [15], using cfDNA collected from the tissue culture media from two lymphoblastoid cell lines from the HapMap/1000 Genome Project, NA12878 (the Platinum Genome sample) and NA11840, to mimic a tumour-normal (or plasma-normal) pair. The dilution series mixed cfDNA from NA12878 with an increasing amount of cfDNA from NA11840 (from 0 to 99.8% by volume, n=12, Additional file 1: Table S1). This cfDNA dilution series was used to investigate the sensitivity in detecting mutations at high and low allele frequency (50% - 0.1%). The 377-amplicon panel encompassed four heterozygous single nucleotide polymorphisms (SNPs) present only in NA12878. These SNPs were used as “somatic” mutations for the purpose of this analysis.

Using NG-TAS the cfDNA dilution series was tested in triplicate, varying the input cfDNA from 5 ng to 50 ng. Since in clinical plasma samples the amount of ctDNA is frequently a limiting factor, we also tested the ThruPlex plasma-seq kit (requiring as little as 3 ng of cfDNA input) to generate a whole genome cfDNA library (termed NGS cfDNA library). An aliquot of this NGS cfDNA library was then used as input for NG-TAS.

These NG-TAS experiments showed a strong linear relationship between the observed and expected variant allele frequencies (VAF) for the four “somatic” mutations (Table 2, Figure 3). As the input cfDNA reduced from 50 ng to 5 ng the R squared values decreased from 0.968 to 0.885. With 10 ng input cfDNA, VAFs as low as 1% could be consistently detected. Lowering the input cfDNA generated more variable results (i.e. VAF deviating from the expected values and higher standard deviations), in particular at low AF. This is probably caused by stochastic amplification of the alternative allele. NG-TAS performed using NGS cfDNA library as input performed better than 5 ng of cfDNA input (R^2^=0.964, Table 2, Figure 3).

**Figure 3.**
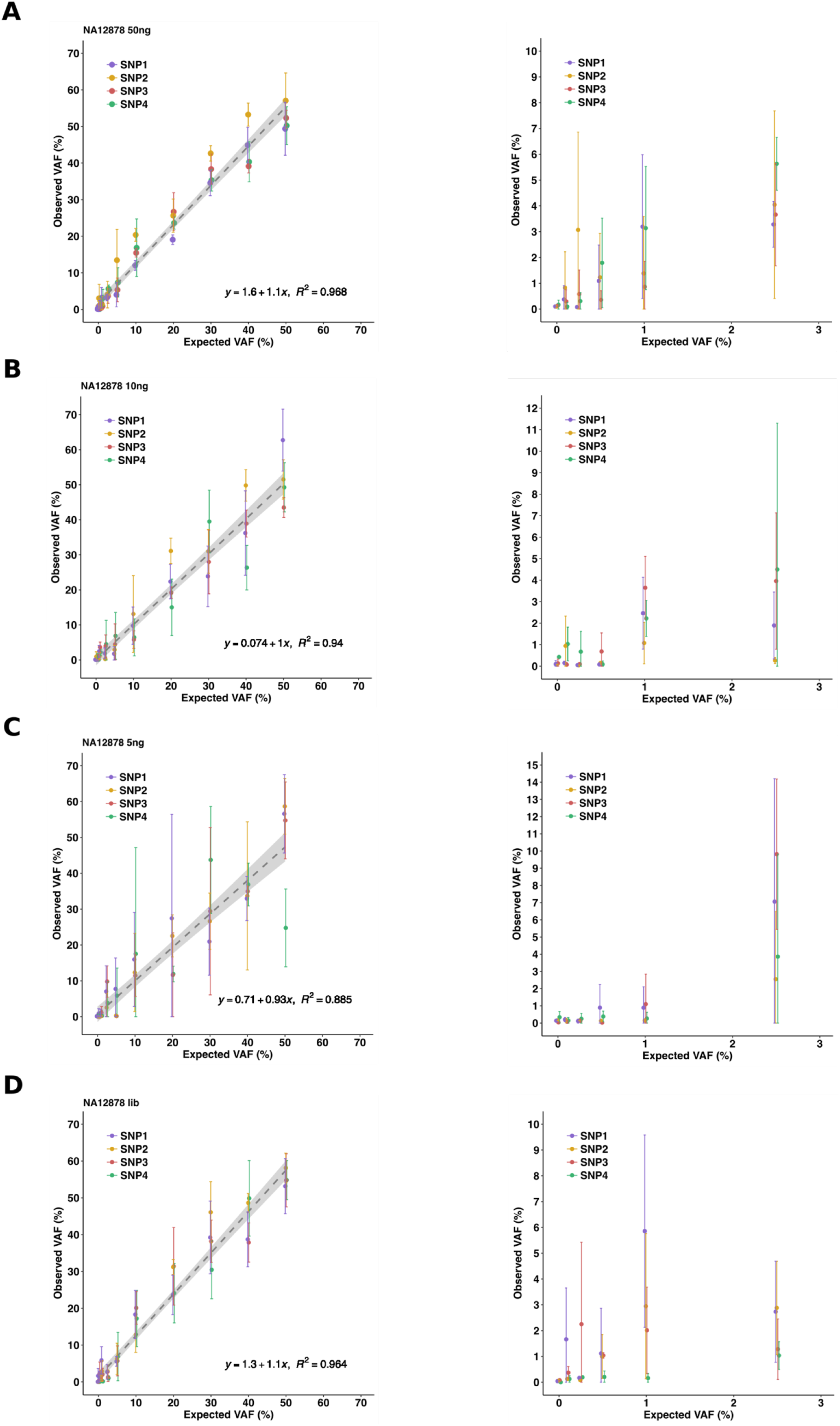
Detection of SNVs in NA12878 cfDNA dilution series. (A) Expected versus observed VAF for 4 SNVs in the NA12878-NA11840 dilution series starting from 50 ng input DNA (left) and zoom-in for expected VAF <5% (right). (B) Expected versus observed VAF for 4 SNVs in the NA12878-NA11840 dilution series starting from 10 ng input DNA (left) and zoom-in for expected VAF <5% (right). (C) Expected versus observed VAF for 4 SNVs in the NA12878-NA11840 dilution series starting from 5 ng input DNA (left) and zoom-in for expected VAF <5% (right). (D) Expected versus observed VAF for 4 SNVs in the NA12878-NA11840 dilution series starting from post-NGS library input DNA (left) and zoom-in for expected VAF <5% (right).

**Table 2.** Linear regression analysis for different cfDNA input.

| Input DNA | R<sup>2</sup> | Estimated coefficient | 2.5% CI | 97.5% CI |
| --- | --- | --- | --- | --- |
| 50ng | 0.968 | 1.075 | 1.018 | 1.133 |
| 10ng | 0.940 | 1.005 | 0.930 | 1.080 |
| 5ng | 0.885 | 0.932 | 0.832 | 1.032 |
| Library | 0.964 | 1.123 | 1.059 | 1.187 |

The NG-TAS analysis pipeline was developed and optimised using this dilution series data and later applied to data from clinical plasma samples. As illustrated in Figure 4A and in the Methods section, mutation calling was performed using MuTect2, processing each amplicon individually. To limit the number of false positives (FPs) caused by PCR errors, we only called mutations observed in at least two out of three replicates. With the reported settings and using 10 ng of input cfDNA from the dilution series, all four SNVs were called when the expected VAF was 5% or higher, and 3 of 4 SNVs when the expected VAF was 1% (Figure 4B). No FPs with VAF higher than 3% were called with 50 ng and 10 ng input cfDNA from the dilution series. NG-TAS of both the 5 ng cfDNA input and NGS cfDNA library input generated seven FPs above 3% in the dilution series (Figure 4C). Template scarcity and extra PCR cycles during library preparation could explain this increase in FPs.

**Figure 4.**
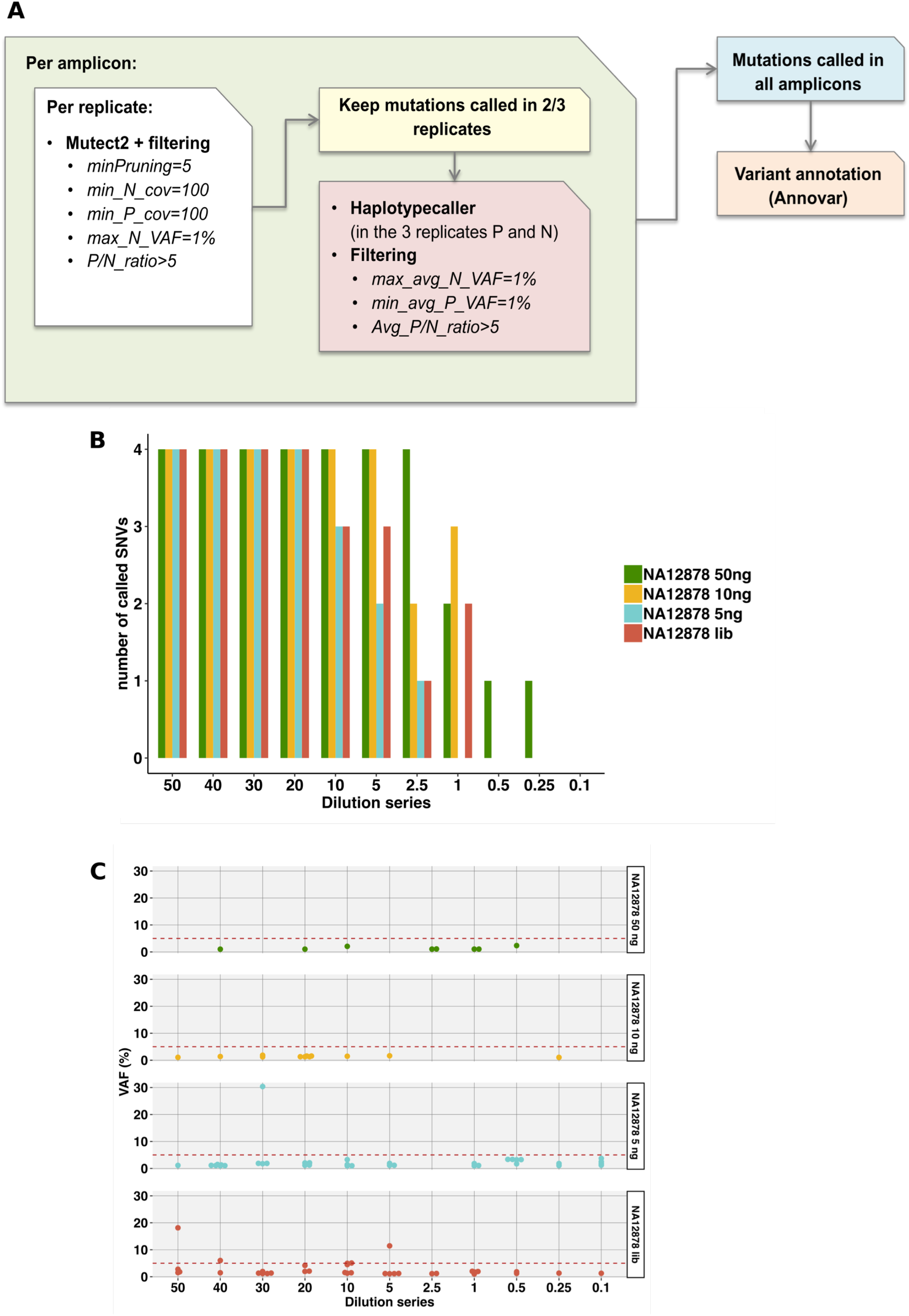
Mutation calling in NA12878 cfDNA dilution series. (A) Schematic overview of the computational pipeline to identify somatic mutations in NG-TAS data. (B) *De novo* mutation calling in the NA12878 dilution series was evaluated for different amounts of input cfDNA. 4 SNVs can potentially be called using our panel of 377 amplicons. (C) VAF for all FP calls in the NA12878 dilution series. The red dashed line represents 5% VAF.

Therefore, for NG-TAS in plasma samples we recommend the use of 10 ng cfDNA per replicate as input, and a threshold of 5% VAF for *de novo* mutation calling. In plasma samples with less cfDNA the use of NGS cfDNA library as input for NG-TAS enables ctDNA profiling in samples with as little as 3 ng of cfDNA. However, this approach is more suitable for tracking in plasma ctDNA mutations previously identified in the tumour, rather than for *de novo* plasma ctDNA mutation calling.

### Testing NG-TAS performance in cancer patient samples

We applied NGTAS to a clinical cohort of 30 metastatic breast cancer patients from which we have collected 360 plasma samples (for 31 of these NGS cfDNA library samples were used) and buffy coats. This cohort is part of a clinical trial which will be comprehensively reported in a separate manuscript (Baird et al, in preparation).

To estimate the FP rate in blood samples, we used pairs of DNA extracted from the buffy coats collected at two different time points from four patients. Any mutation identified by NG-TAS in any of the eight possible buffy coat DNA pairs tested was considered a FP. Figure 5A shows that in these samples NG-TAS identified no FP with VAF greater than 5% (a result similar to NG-TAS performed using the cell line cfDNA dilution series, Figure 4C).

**Figure 5.**
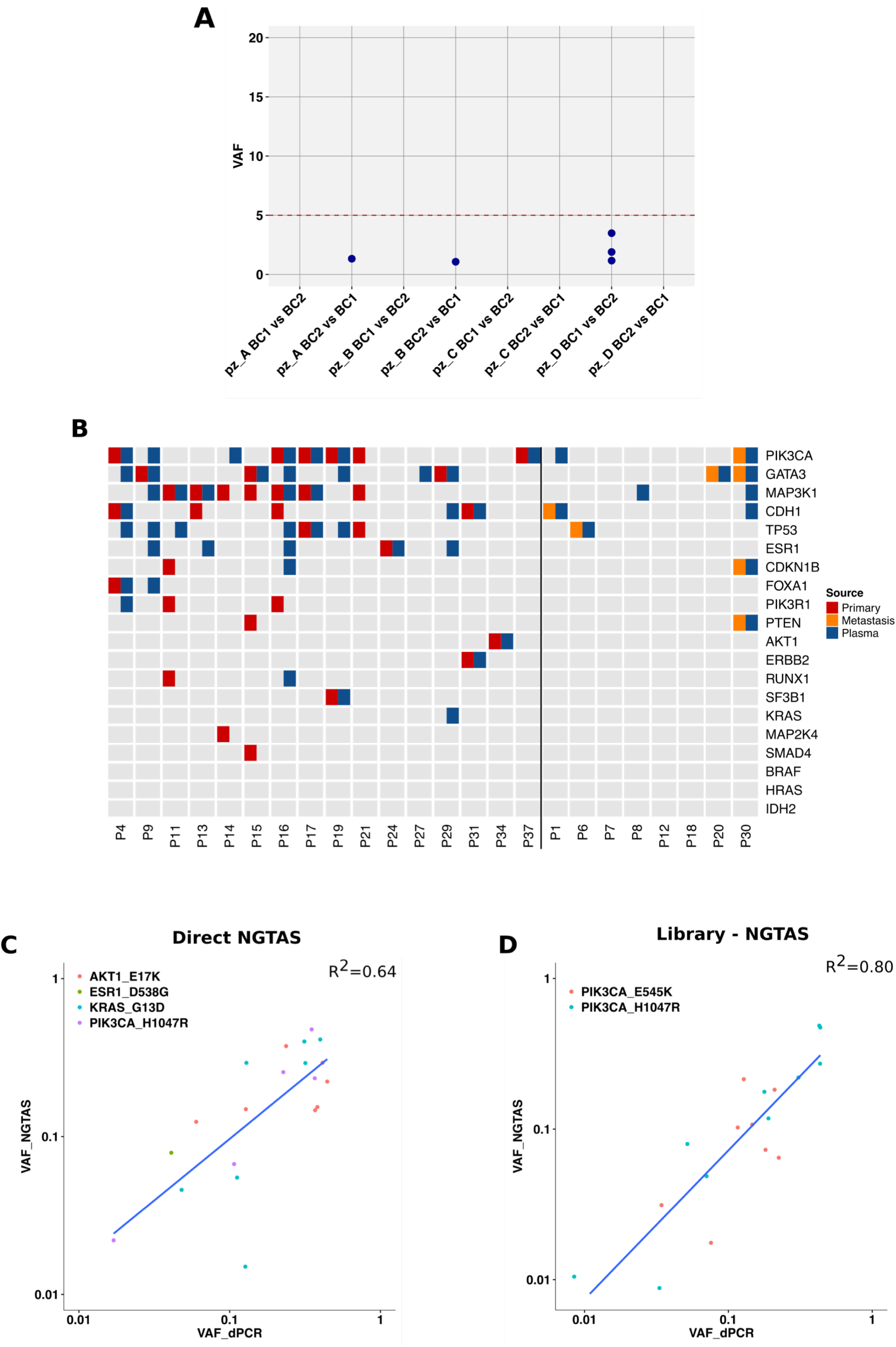
Validation of NG-TAS performance in clinical plasma samples. (A) The specificity of NG-TAS in clinical samples was estimated using 4 pairs of buffy coats from the same patients (A, B, C and D). The mutation calling pipeline was applied using one buffy coat as normal and the other as ‘tumour’ and vice versa. All mutations called in this setting can be considered FPs. The red line indicates 5% VAF. (B) Oncoprint summary plot of genes mutated in 24 cases for which both tissue and plasma samples were tested. The vertical black line separates cases for which the primary tumour was analysed from cases for which a metastasis biopsy was analysed. (C-D) Comparison of VAF obtained by NG-TAS and dPCR. (C) In this comparison, four different hotspot mutations including *AKT1* (*E17K*), *ESR1* (*D538G*), *KRAS* (*G13D*) and *PIK3CA* (*H1047R*) identified in multiple plasma samples from 4 distinct patients were analysed (R^2^ = 0.64). (D) Two *PIK3CA* hotspots (*H1047R* and *E545K*) were detected by NG-TAS using NGS library as input material in plasma samples from two distinct patients. The same mutations were detected using dPCR and a good correlation was found (R^2^ = 0.80).

In 24 of the cases in our cohort, at least one tissue sample was also available and analysed. Sixteen of these cases had tissues from the primary tumour while in the remaining 8 cases, tissue samples were obtained from metastasis biopsies collected during the trial. Overall, we found at least one mutation in 21/24 patients (87.5%, Figure 5B). Forty-four mutations were detected in the tissue samples and 60 in at least one plasma sample; of these, 23 were observed in both tissue and plasma. The agreement was higher for the 8 cases where a metastasis biopsy was sequenced: 7 mutations detected in the tissue, 11 detected in plasma, 7 in common (100% of tissue mutations detected in plasma). In the 16 cases where a primary tumour was tested, 33 mutations were detected in the tissue, 41 in plasma, 19 in common (58% of tissue mutations detected in plasma, Figure 5B and Additional file 1: Fig. S5). The discordance seen in this cohort is probably due to the time gap between the primary tumour tissue sample and plasma, the latter obtained when the patients had metastatic disease. In addition, most of the tissue samples were formalin-fixed paraffin-embedded (FFPE) hence, we detected an increase of C>T/G>A SNVs not usually found in ctDNA samples (Additional file 1: Fig. S5).

We used dPCR to validate a subset of the mutations identified in seven patients in which NG-TAS was performed either directly on cfDNA (n=4) or using post-NGS library products (n=3). In the four direct NG-TAS samples, four hotspot mutations *PIK3CA* (*H1047R* and *E545K*), *KRAS* (*G13D*), *ESR1* (*D538G*) and *AKT1* (*E17K*)) were all validated by dPCR. A good concordance between VAFs estimated by NG-TAS and dPCR was found (R^2^=0.64, Figure 5C). In the three patients where post-NGS library products was used as input, two *PIK3CA* hotspots (*H1047R and E545K*) were also validated by dPCR and a high concordance between the VAFs estimated by NG-TAS and dPCR was observed (R^2^=0.80, Figure 5D).

### Monitoring response in breast cancer patients using NG-TAS

We report the example of two patients from the above clinical trial to demonstrate the use of NG-TAS for metastatic breast cancer disease monitoring. Patients had clinical monitoring performed as per the trial protocol using RECIST (Response Evaluation Criteria in Solid Tumour), version 1.1.

The first patient had RECIST partial response in the first 28 weeks, and progression on day 197. NG-TAS identified mutations in *GATA3* (*F431fs*), *PIK3CA* (*E542K*), *CDKN1B* (*N124fs*) and *PTEN* (*137-139del*) (Figure 6A). *PTEN* mutation VAFs in ctDNA showed parallel dynamics to RECIST: initial drop, followed by continuous rise from day 85, preceding RECIST progression by over 100 days. The VAFs of the other mutations showed a parallel rise starting later.

**Figure 6.**
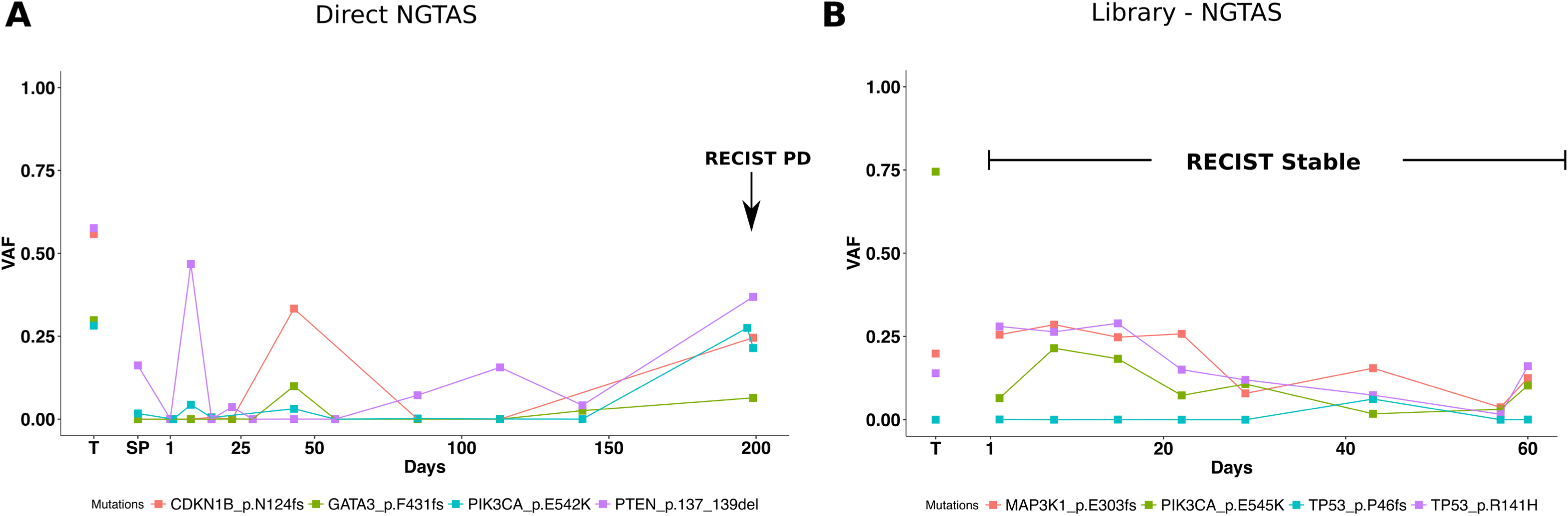
Monitoring response in metastatic breast cancer patients using NG-TAS. (A) Example of patient monitoring during treatment using direct NG-TAS in ctDNA. There are four mutations detected in more than one sample: *GATA3* (*F431fs*), *PIK3CA* (*E542K*), *CDKN1B* (*N124fs*) and *PTEN* (*137-139del*). The mutations called more than once in the longitudinal samples are shown including tumour and plasma samples. The arrow indicates the time of the disease considered as RECIST progressive disease. T indicates tumour samples, and SP indicates screening plasma sample which was collected prior to the treatment. (B) Example of patient monitoring during treatment using NGS library material for NG-TAS. This patient had a stable disease during the whole treatment period. There are three mutations detected, including *MAP3K1* (*E303 frame shift*), hotspot mutations *PIK3CA* (*E545K*) and *TP53* (*R141H* and *P46fs*). T indicates tumour samples.

The second patient had stable disease by RECIST during the 60 days of available follow up. Due to limited amount of cfDNA extracted in this case, NG-TAS was performed using NGS cfDNA libraries. NG-TAS detected *PIK3CA* (*H1047R*), *MAP3K1* (*E303fs*) and *TP53* (*R141H* and *P46fs*) mutations, and their VAFs showed stable values, then a slight reduction between days 20-56, followed by a slightly rise by the time monitoring was discontinued (Figure 6B).

These two examples demonstrate the use of NG-TAS in plasma cfDNA samples to monitor tumour burden in metastatic breast cancer patients.

### Comparison of NG-TAS with other approaches

We finally compared NG-TAS to other existing technologies such as digital PCR, TAm-Seq and Oncomine™ Breast cfDNA Assay (Table 3). NG-TAS can be performed in 7 hours using the Fluidigm system as detailed in the methods session. Up to 384 samples can be processed at the same time. Lower limits of detection can be reached using Digital PCR or Oncomine technology, however this is limited to one target for the first and a set of pre-defined hotspots for the latter. Importantly, the cost of NG-TAS, estimated at 30 GBP per sample is significantly lower than any commercial solution, making it cost-effective for use in the clinics.

**Table 3.**
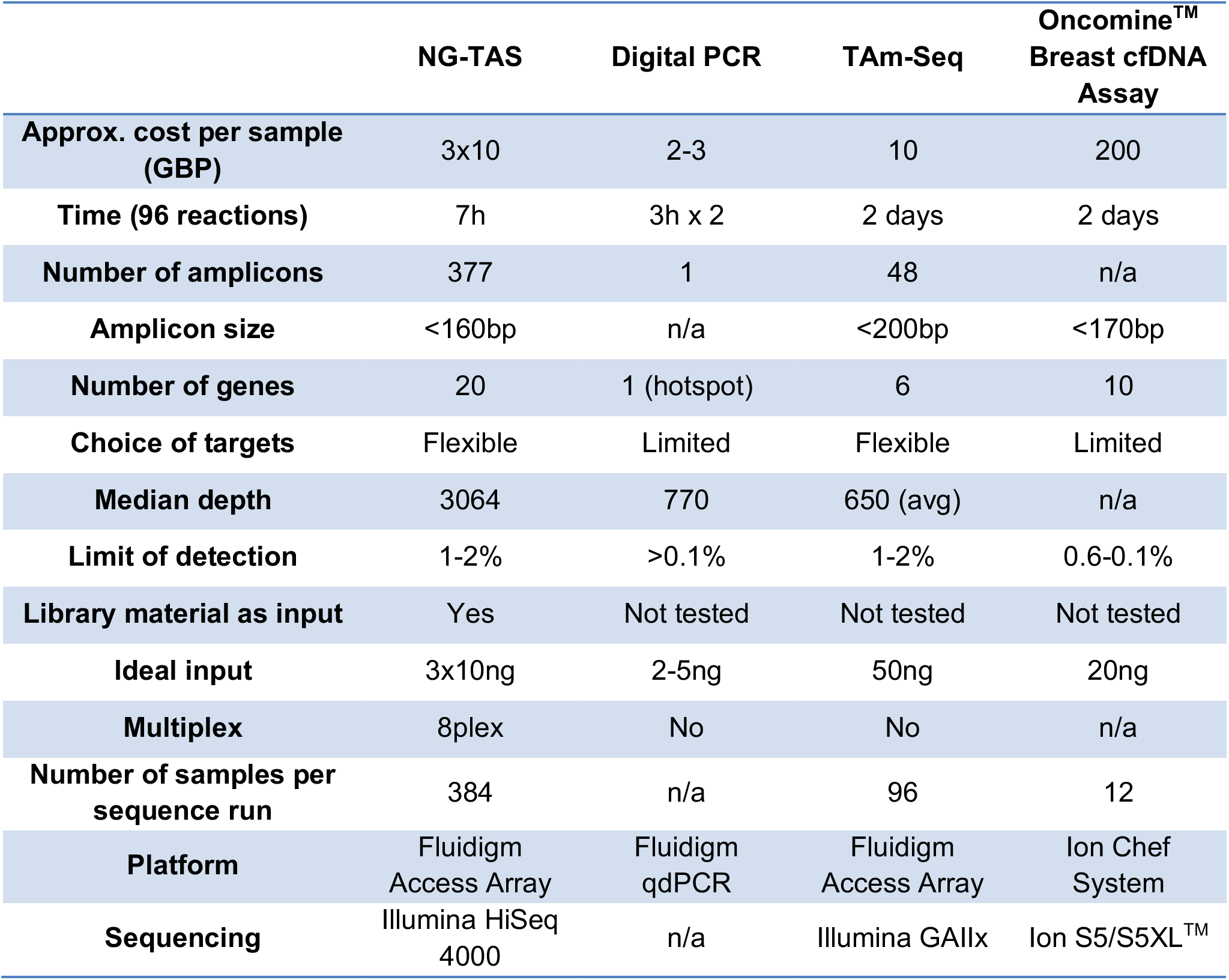
Comparison of different approaches for ctDNA detection.

|  | NG-TAS | Digital PCR | TAm-Seq | Oncomine™<br>Breast cfDNA<br>Assay |
| --- | --- | --- | --- | --- |
| **Approx. cost per sample (GBP)** | 3x10 | 2-3 | 10 | 200 |
| **Time (96 reactions)** | 7h | 3h x 2 | 2 days | 2 days |
| **Number of amplicons** | 377 | 1 | 48 | n/a |
| **Amplicon size** | <160bp | n/a | <200bp | <170bp |
| **Number of genes** | 20 | 1 (hotspot) | 6 | 10 |
| **Choice of targets** | Flexible | Limited | Flexible | Limited |
| **Median depth** | 3064 | 770 | 650 (avg) | n/a |
| **Limit of detection** | 1-2% | >0.1% | 1-2% | 0.6-0.1% |
| **Library material as input** | Yes | Not tested | Not tested | Not tested |
| **Ideal input** | 3x10ng | 2-5ng | 50ng | 20ng |
| **Multiplex** | 8plex | No | No | n/a |
| **Number of samples per sequence run** | 384 | n/a | 96 | 12 |
| **Platform** | Fluidigm<br>Access Array | Fluidigm<br>qdPCR | Fluidigm<br>Access Array | Ion Chef<br>System |
| **Sequencing** | Illumina HiSeq<br>4000 | n/a | Illumina GAIIx | Ion S5/S5XL™ |

## DISCUSSION

The genes frequently mutated in different human cancers have been characterized by large-scale sequencing studies such as The Cancer Genome Atlas [20,21]. These pan-cancer studies have revealed that most human tumours have at least 1-10 driver mutations, allowing the design of custom gene panels that could be used for generic cancer detection. But the challenge remaining is there are very few recurrent or hotspot mutations in tumours such as breast cancer, with mutations spread along the protein coding region, as observed in *TP53, GATA3* and *MAP3K1.* Therefore, it would be desirable to cover most exons of these genes simultaneously in a ctDNA mutation detection panel.

The detection of specific mutations in ctDNA is achievable by dPCR, now considered the gold standard to detect mutations with low VAFs. However, dPCR is constrained by the number of mutations that can be detected in a single reaction [11]. Thus, its high sensitivity and specificity is at the expense of the number of mutations that can be detected concurrently. At the other end of the spectrum, whole genome sequencing or whole exome sequencing suffer from reduced sensitivity at the current achievable level of sequencing depth [22].

We report here a new approach, NG-TAS, an optimised targeted amplicon sequencing pipeline that provides clinically relevant sensitivity in mutation calling across a targeted, but relatively broad and customizable panel of genes. The current version of NG-TAS covers all exons or hotspots of 20 breast cancer-associated genes in a total of 377 amplicons, has a lower detection limit of 1% VAF, and requires only three aliquots of 10 ng cfDNA input. The single step multiplexed PCR amplification makes it a less time consuming method and more cost effective than other assays, such as the commercially available Oncomine assay (Table 3). NG-TAS is flexible and custom designed primers can be adjusted to the needs of the end user, depending on the cancer type and the clinical context.

Importantly, we developed a bespoke NG-TAS computational pipeline for data analysis, with all the relevant open-source code available at GitHub (https://github.com/cclab-brca/NGTAS_pipeline). All sequencing data are also made available at: https://figshare.com/articles/NGTASNA12878/7387370 and https://www.ebi.ac.uk (EGAS00001003392). These will be instrumental to test and further develop the computational pipeline, as required by regulatory agencies.

The custom design of primers for NG-TAS is potentially challenging. Building a customised panel of primers manually, using the tool mentioned above is time-consuming and, in some cases difficult due to genomic sequence context (e.g. high GC and repetitive regions). The multiplex PCR requires a fixed annealing temperature, but more complex PCR cycle design can circumvent this. Nevertheless, we were able to design primers that yielded in 94% of amplicons over 100x coverage (Figure 2B). We provide all primer sequences (Additional file 2) and an open source optimised primer library will be growing with an NG-TAS user community.

When using NG-TAS for accurate estimation of VAF, as required to do serial tumour burden monitoring, our data suggests that at least 10 ng of input cfDNA per replicate is required. NG-TAS has poor performance with cfDNA input below 5 ng (per replicate), with amplicon coverage reduced in a stochastic manner, probably due to the limited availability of template. A suitable alternative protocol for these cases is to generate an NGS cfDNA library, requiring only 3 ng of cfDNA, and use the library material as input for NG-TAS.

We applied NG-TAS to a cohort of 30 patients for which both tissue and serial plasma samples were available. The percentage of mutations identified in tissue and detected in ctDNA was 100% when the tissue was from a synchronous metastasis biopsy and 58% when the tissue was from the primary tumour. Such agreement is higher than what recently reported by Chae et al [23]. In their cohort of 45 patients, 60% of tissue samples were from primary tumours and 58% of the tissues were acquired more than 90 days before ctDNA testing. The FoundationOne panel was used for tissue analysis and the Guardant360 assay for ctDNA. They detected only 25.6% of tissue mutations in plasma when evaluating the common regions between the two targeted approaches.

A future development of NG-TAS will be the use of molecular barcoding, since this has been shown to improve sensitivity and specificity of amplicon-based deep sequencing [24]. This will have cost implications, potentially limiting one of the main advantages of the current NG-TAS protocol. The extra costs would be the result of generation of barcoded primers. For example, if 96 distinct barcodes are used, the primer cost will increase around 100 times. However, costs will be significantly diluted when considering laboratories processing a large number of samples, keeping the overall cost of NG-TAS within a very reasonable range.

## Conclusions

We have described here the workflow for a highly multiplexed cfDNA deep sequencing method named NG-TAS. NG-TAS assesses the mutational status of several genes simultaneously, with high sensitivity (allowing quantification of AF) and competitive costs, and offers flexibility in the choice of target genes. We have also shown proof of principle that the monitoring of ctDNA using NG-TAS in metastatic breast cancer can allow detection of cancer progression earlier than conventional RECIST measurements.

## List of abbreviations

NG-TAS: Next Generation Targeted Amplicon Sequencing
cfDNA: Cell Free DNA
ctDNA: Circulating Tumour DNA
dPCR: Digital PCR
TAm-Seq: Targeted Amplicon Sequencing
NGS: Next Generation Sequencing
FFPE: Formalin Fixed Paraffin Embedded
VAF: Variant Allele frequency
SNV: Single Nucleotide Variant
FP: False Positive
UDG: Uracil DNA Glycosylase
RECIST: Response Evaluation Criteria in Solid Tumour
CT: Computed Tomography

## Declarations

### Ethics approval and consent to participate

This study was approved by the East of England - Cambridge East Research Ethics Committee (REC reference: 14/EE/1045). All human samples used were collected after informed consent, and the study was fully compliant with the Helsinki Declaration.

## Consent for publication

Not applicable

## Availability of data and material

The computational pipeline is available through GitHub (https://github.com/cclab-brca/NGTAS_pipeline). All sequencing data are available at: https://figshare.com/articles/NGTAS_NA12878/7387370 and https://www.ebi.ac.uk (EGAS00001003392).

## Authors’ contributions

MG, MC, EB, SFC and CC conceived the study; MG and EB designed primers and generated the dilution series; MG performed NG-TAS in Fluidigm Access Array™ system; RB, KB, SCL, JR, JC and MO conducted the clinical trial; HB, LJ and AB collected the clinical samples; MC developed the computational approach and performed the analyses; SJS contributed in the computational pipeline development; MC and MG performed data analysis; MG, MC, SFC and CC drafted the manuscript, all authors revised and approved the final manuscript.

## Acknowledgements

We are grateful to Cancer Research UK and the University of Cambridge for the support. We thank the Cancer Molecular Diagnostics Lab and Cancer Research UK Cambridge Institute Core Facilities including Genomics and Bio-repository that supported this work. We thank Dr PA Edwards for the scientific advice and editing for this manuscript.

## Additional files

**Additional file 1:** Supplementary figures and tables, PDF 722 KB

**Additional file 2:** List and description of the 377 primers used, XLS 214 KB

