## Additional file 1 for "Next Generation-Targeted Amplicon Sequencing (NG-TAS): An optimised protocol and computational pipeline for cost-effective profiling of circulating tumour DNA"

**Supplementary Figures**

**Fig. S1** – Schematic overview of the computational pipeline to identify somatic mutations in NG-TAS data from longitudinal samples.

**Fig. S2** - Representative image of the Bioanalyser gel plot. The 8plex PCR products were analysed using Bioanalyser for primer efficiency and quality control.

**Fig. S3** - Fragment size distribution according to the Bioanalyser results for cfDNA extracted from the media where NA12878 cells were grown (main peak at around 160-170bp).

**Fig. S4** – (A) Percentage of amplicons having more than 100x coverage for 2, 5 and 10 ng of input cfDNA from NA12878 sample. (B) Percentage of reads on target for 2, 5 and 10 ng of input cfDNA from NA12878 sample.

**Fig. S5** – Detailed representation of mutations identified in tumour or plasma samples of 21 metastatic breast cancer cases. The colour gradient indicates the VAF as indicated; PT = primary tumour, M = metastasis biopsy, V1…n = plasma.

### Fig S1

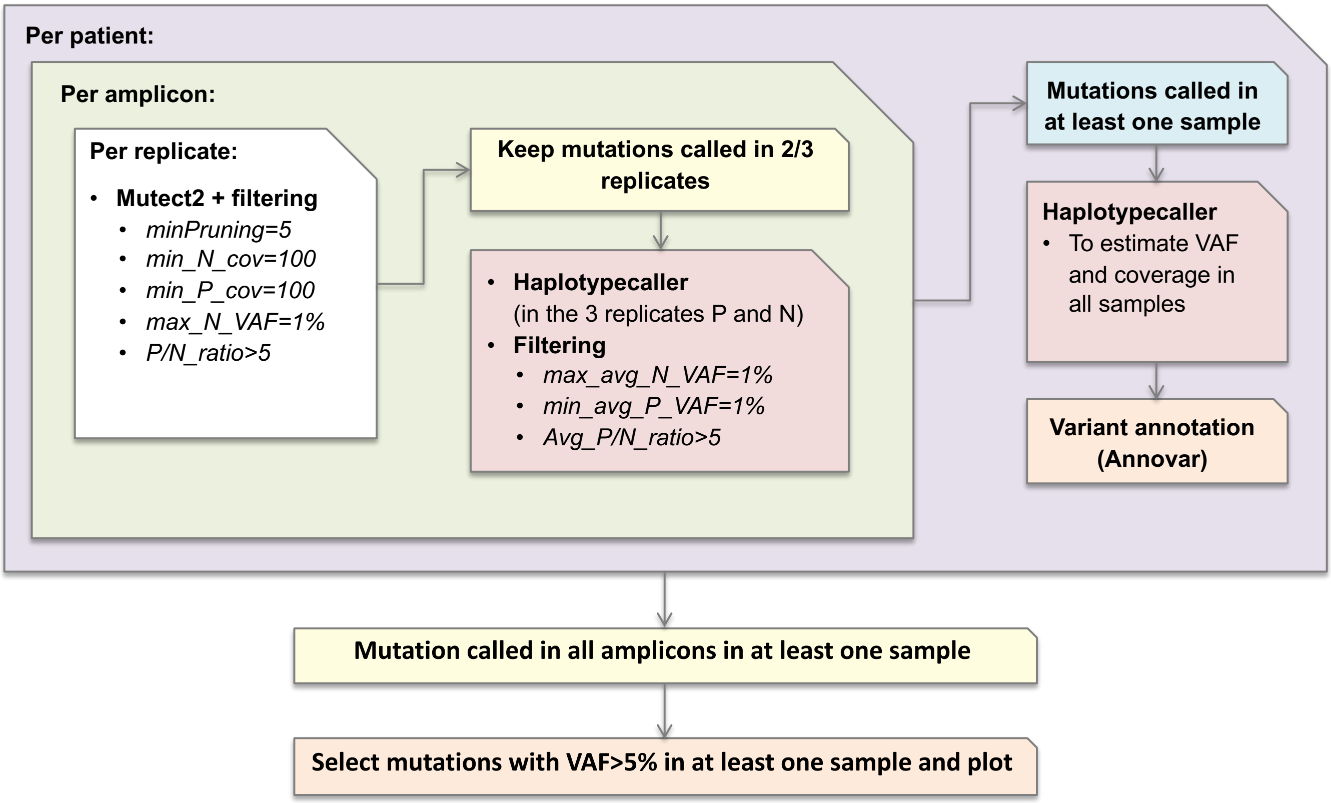
Per patient:
Per amplicon:
Per replicate:
Mutect2 + filtering
minPruning=5
min_N_cov=100
min_P_cov=100
max_N_VAF=1%
P/N_ratio>5
Keep mutations called in 2/3 replicates
Haplotypecaller
(in the 3 replicates P and N)
Filtering
max_avg_N_VAF=1%
min_avg_P_VAF=1%
Avg_P/N_ratio>5
Mutations called in at least one sample
Haplotypecaller
To estimate VAF and coverage in all samples
Variant annotation (Annovar)
Mutation called in all amplicons in at least one sample
Select mutations with VAF>5% in at least one sample and plot

### Fig S2

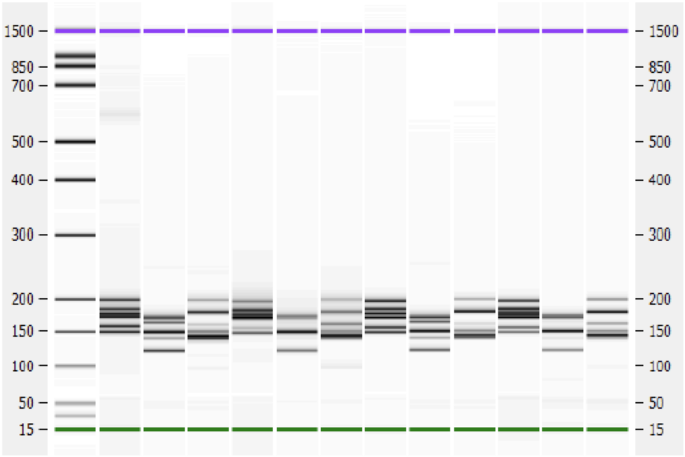
1500
850
700
500
400
300
200
150
100
50
15

### Fig S3

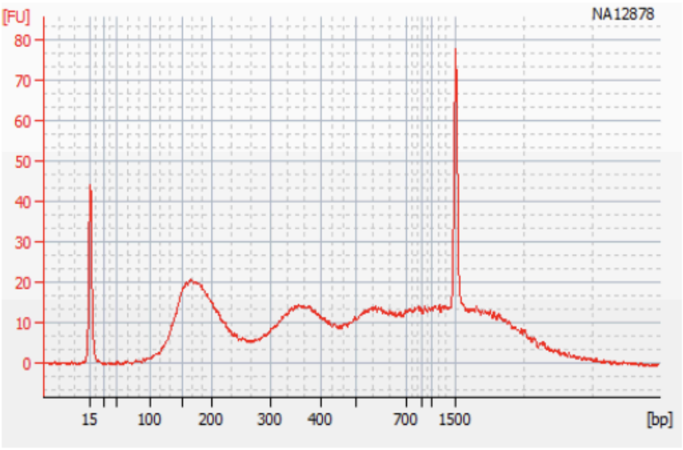
[FU]
NA12878
80
70
60
50
40
30
20
10
0
15
100
200
300
400
700
1500
[bp]

**Fig S4**

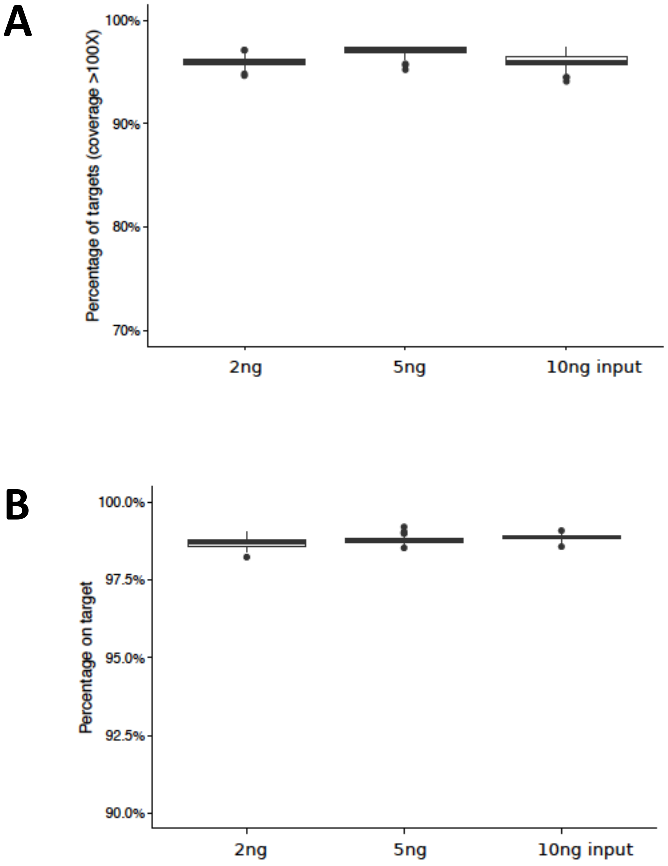
A
100%
90%
80%
70%
Percentage of targets (coverage >100X)
2ng
5ng
10ng input
B
100.0%
97.5%
95.0%
92.5%
90.0%
Percentage on target
2ng
5ng
10ng input

### Fig S5

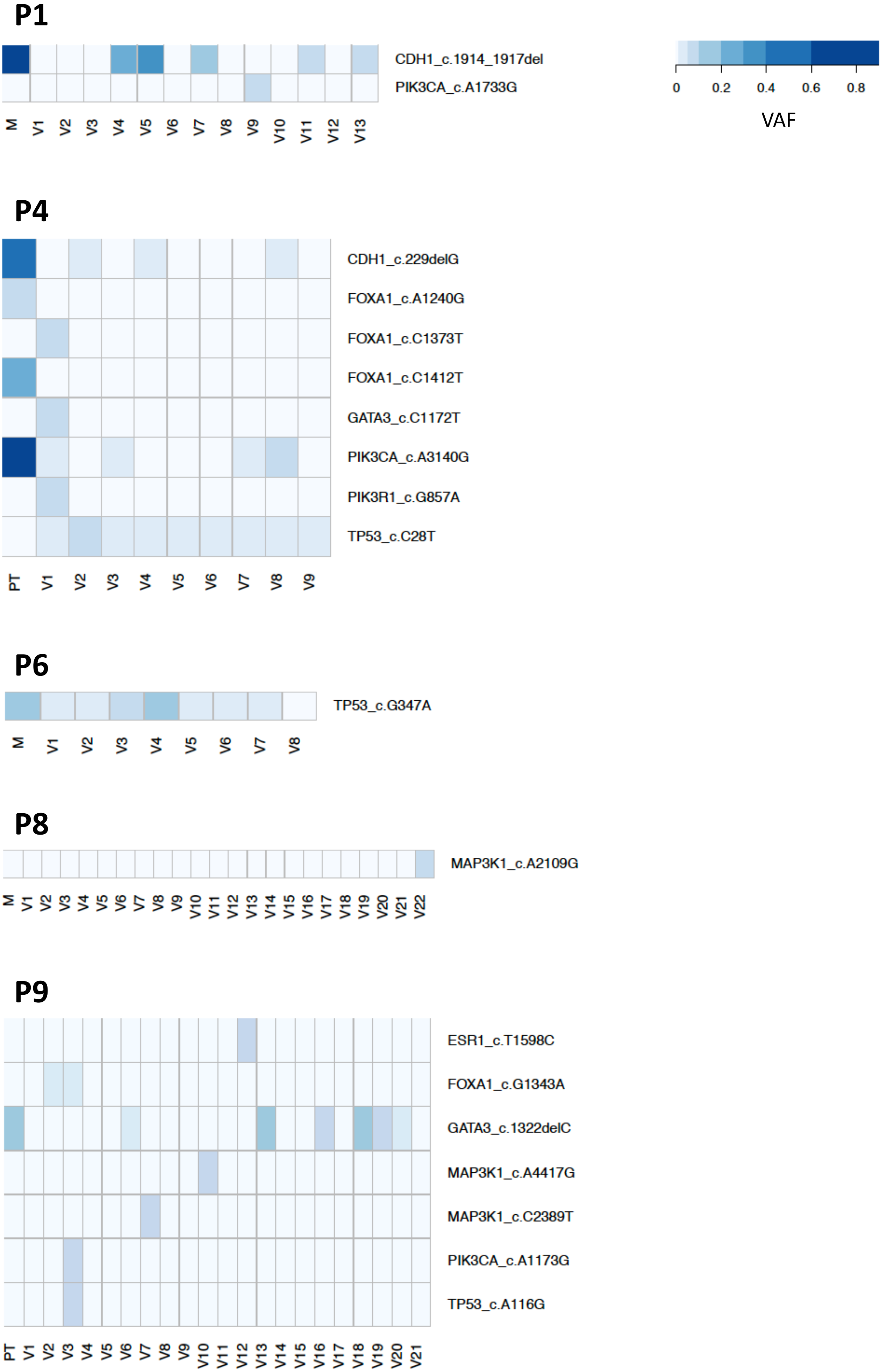

P1
CDH1_c.1914_1917del
PIK3CA_c.A1733G
M V1 V2 V3 V4 V5 V6 V7 V8 V9 V10 V11 V12 V13
0 0.2 0.4 0.6 0.8
VAF
P4
CDH1_c.229delG
FOXA1_c.A1240G
FOXA1_c.C1373T
FOXA1_c.C1412T
GATA3_c.C1172T
PIK3CA_c.A3140G
PIK3R1_c.G857A
TP53_c.C28T
PT V1 V2 V3 V4 V5 V6 V7 V8 V9
P6
TP53_c.G347A
M V1 V2 V3 V4 V5 V6 V7 V8
P8
MAP3K1_c.A2109G
M V1 V2 V3 V4 V5 V6 V7 V8 V9 V10 V11 V12 V13 V14 V15 V16 V17 V18 V19 V20 V21 V22
P9
ESR1_c.T1598C
FOXA1_c.G1343A
GATA3_c.1322delC
MAP3K1_c.A4417G
MAP3K1_c.C2389T
PIK3CA_c.A1173G
TP53_c.A116G
PT V1 V2 V3 V4 V5 V6 V7 V8 V9 V10 V11 V12 V13 V14 V15 V16 V17 V18 V19 V20 V21

## P11

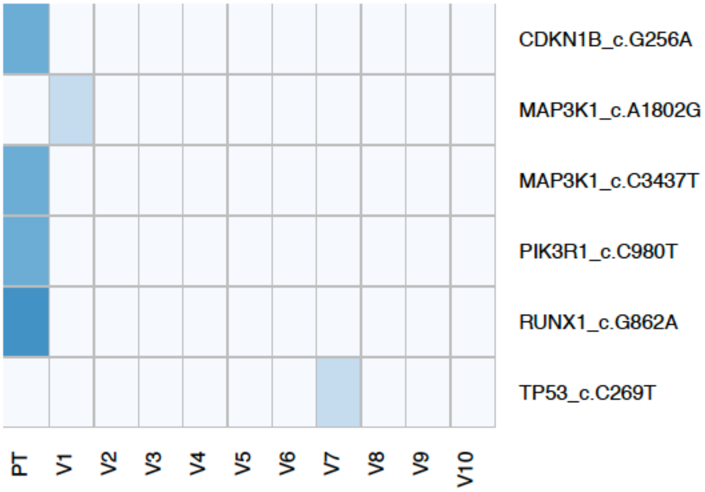
CDKN1B_c.G256A
MAP3K1_c.A1802G
MAP3K1_c.C3437T
PIK3R1_c.C980T
RUNX1_c.G862A
TP53_c.C269T
PT
V1
V2
V3
V4
V5
V6
V7
V8
V9
V10

## P13

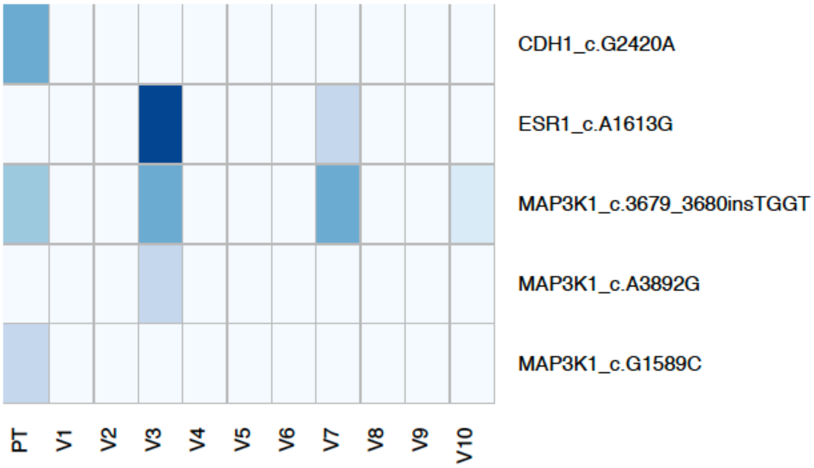
CDH1_c.G2420A
ESR1_c.A1613G
MAP3K1_c.3679_3680insTGGT
MAP3K1_c.A3892G
MAP3K1_c.G1589C
PT
V1
V2
V3
V4
V5
V6
V7
V8
V9
V10

## P14

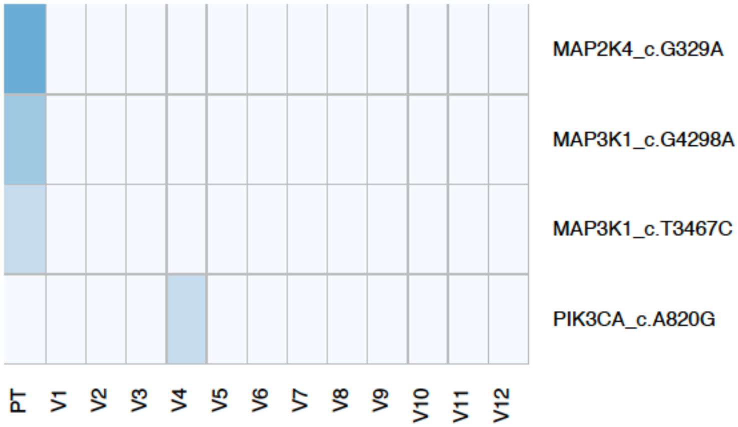
MAP2K4_c.G329A
MAP3K1_c.G4298A
MAP3K1_c.T3467C
PIK3CA_c.A820G
PT
V1
V2
V3
V4
V5
V6
V7
V8
V9
V10
V11
V12

**P15**

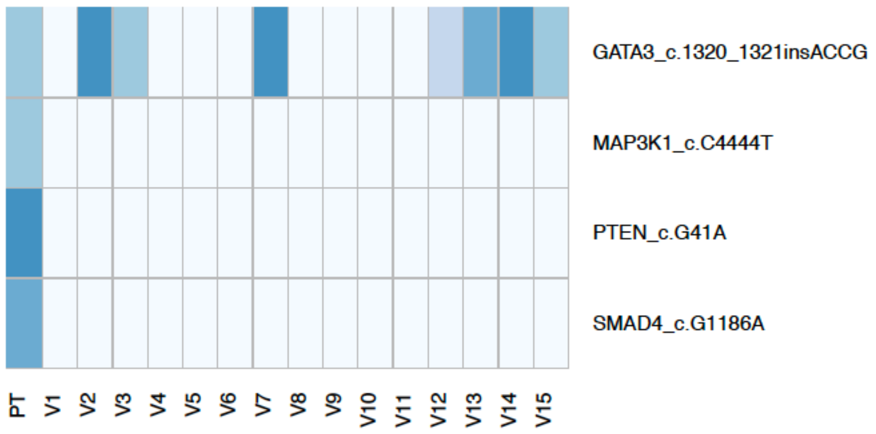
GATA3_c.1320_1321insACCG
MAP3K1_c.C4444T
PTEN_c.G41A
SMAD4_c.G1186A
PT
V1
V2
V3
V4
V5
V6
V7
V8
V9
V10
V11
V12
V13
V14
V15

**P16**

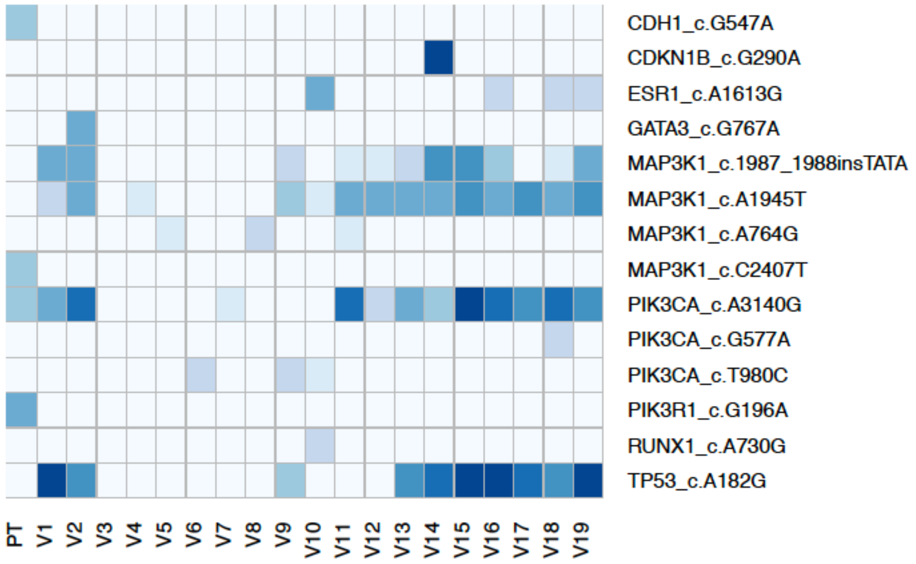
CDH1_c.G547A
CDKN1B_c.G290A
ESR1_c.A1613G
GATA3_c.G767A
MAP3K1_c.1987_1988insTATA
MAP3K1_c.A1945T
MAP3K1_c.A764G
MAP3K1_c.C2407T
PIK3CA_c.A3140G
PIK3CA_c.G577A
PIK3CA_c.T980C
PIK3R1_c.G196A
RUNX1_c.A730G
TP53_c.A182G
PT
V1
V2
V3
V4
V5
V6
V7
V8
V9
V10
V11
V12
V13
V14
V15
V16
V17
V18
V19

**P17**

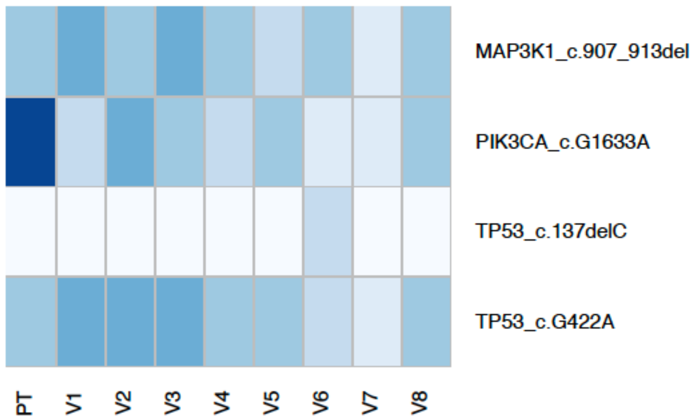
MAP3K1_c.907_913del
PIK3CA_c.G1633A
TP53_c.137delC
TP53_c.G422A
PT
V1
V2
V3
V4
V5
V6
V7
V8

## P19

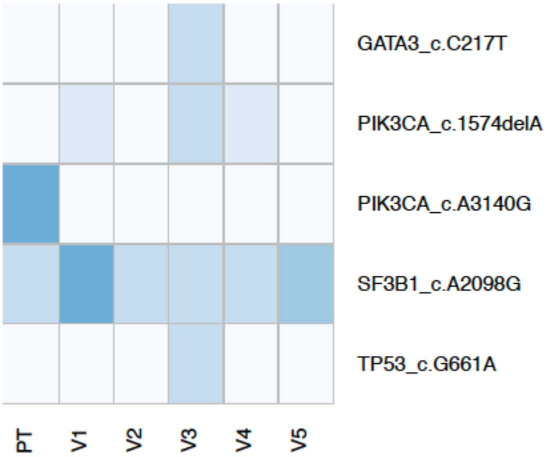


## P20

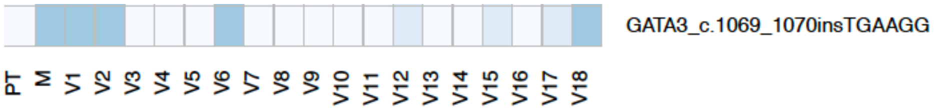


## P21

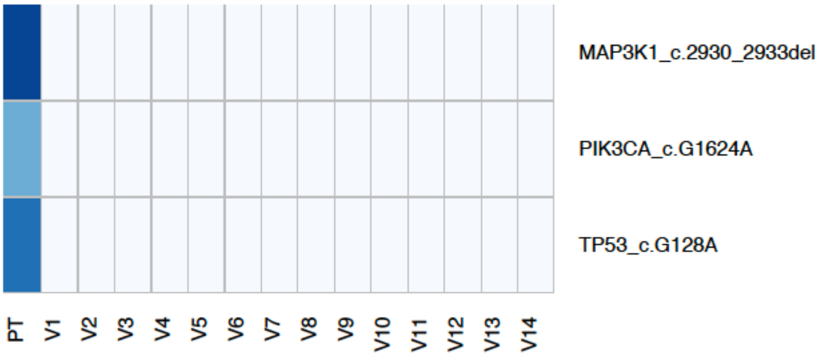


## P24

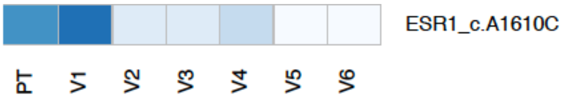


## P27

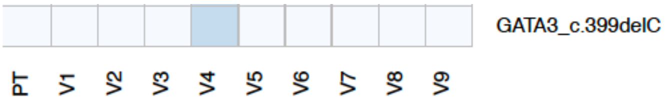

## P29

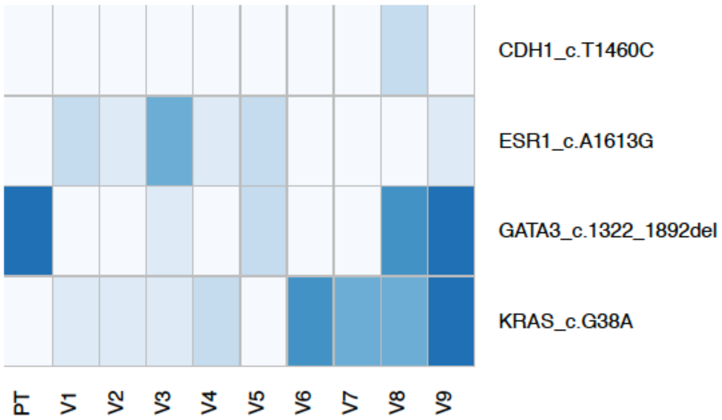


## P30

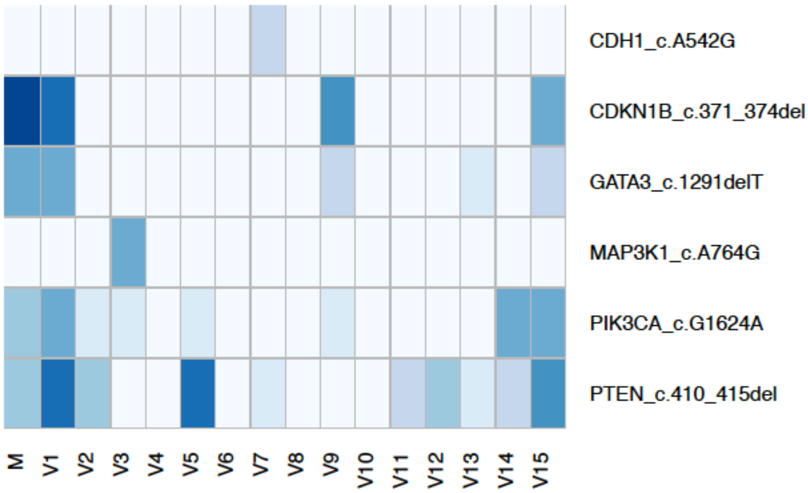


## P31

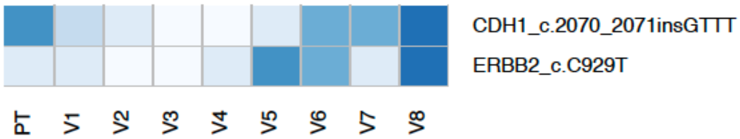


## P34

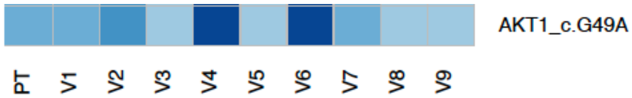


## P37

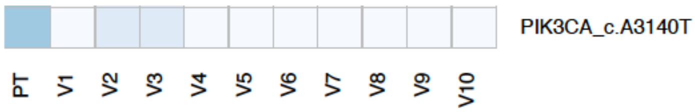

### Table S1: The proportion of NA12878 and NA11840 for the generation of the cfDNA dilution series with expected VAF

| Expected VAF (%) | NA12878 | NA11840 |
|---|---|---|
| **50** | 100 | 0 |
| **40** | 80 | 20 |
| **30** | 60 | 40 |
| **20** | 40 | 60 |
| **10** | 20 | 80 |
| **5** | 10 | 90 |
| **2.5** | 5 | 95 |
| **1** | 2 | 98 |
| **0.5** | 1 | 99 |
| **0.25** | 0.5 | 99.5 |
| **0.1** | 0.2 | 99.8 |
| **0** | 0 | 100 |

### Table S2: Primers and Probes for PIK3CA and ESR1 hotspot mutations for digital PCR

| Gene | Assay | F Primer | R Primer | WT probe (5' VIC/3' MGB) | Mutant probe (5' 6-FAM/3' MGB) |
|---|---|---|---|---|---|
| PIK3CA | p.E545K (c.1633 G>A) | GCAATTTCTACACGAGATCCTCTCT | CATTTTAGCACTTACCTGTGACTCCAT | TGAAATCACTGAGCAGGAG | TGAAATCACTAAGCAGGA |
| | p.H1047R (c.3140 A>G) | AAGAGGCTTTGGAGTATTTCATGAA | TGTTTAATTGTGTGGAAGATCCAATC | CAAATGAATGATGCACATC | TGATGCACGTCATGGT |
| ESR1 | p.D538G (c.1613 A>G) | AGGCATGGAGCATCTGTACA | TTGGTCCGTCTCCTCCA | TGGTGCCCCTCTATGACCTG | CCCTCTATGGCCTGCTGCT |
